# No support for the sexy-sperm hypothesis in the seed beetle: sons of monandrous females fare better in post-copulatory competition

**DOI:** 10.1101/336883

**Authors:** Kristin A. Hook

## Abstract

The sexy-sperm hypothesis posits that polyandrous females derive an indirect fitness benefit from multi-male mating because they increase the probability their eggs are fertilized by males whose sperm have high fertilizing efficiency, which is assumed to be heritable and conferred on their sons. However, whether this process occurs is contentious because father-to-son heritability may be constrained by the genetic architecture underlying traits important in sperm competition within certain species. Previous empirical work has revealed such genetic constraints in the seed beetle, *Callosobruchus maculatus*, a model system in sperm competition studies in which female multi-male mating is ubiquitous. Using the seed beetle, I tested a critical prediction of the sexy-sperm hypothesis that polyandrous females produce sons that are on average more successful under sperm competition than sons from monandrous females. Contrary to the prediction of the sexy-sperm hypothesis, I found that sons from monandrous females had significantly higher relative paternity in competitive double matings. Moreover, post-hoc analyses revealed that these sons produced significantly larger ejaculates when second to mate, despite being smaller. This study is the first to provide empirical evidence for post-copulatory processes favoring monandrous sons and discusses potential explanations for the unexpected bias in paternity.

## INTRODUCTION

Polyandry is a taxonomically widespread mating system whereby a female mates with two or more males over a single reproductive cycle (Thornhill and Alcock 1983). Given that such a phenomenon likely has associated costs and that often a single mating results in more sperm than is required for fertilizing ova, why females mate with multiple males remains an evolutionary enigma (Slatyer et al. 2012). Multiple adaptive hypotheses exist for polyandry and may be divided into those in which the female directly benefits through her own enhanced fitness (i.e., survival and/or reproductive success) and those in which the female indirectly benefits through the enhanced fitness of her offspring (Jennions and Petri 2000; Birkhead and Pizzari 2002). One hypothesis that incites indirect fitness benefits for female multi-male mating is the “sexy-sperm” hypothesis, which posits that polyandrous females increase the probability their eggs are fertilized by males whose sperm have high fertilizing efficiency, which is assumed to be heritable and thus conferred on their sons (Curtsinger 1991; Keller and Reeve 1995).

Analogous to Fisherian runaway selection (‘sexy sons’), the sexy-sperm hypothesis suggests that the female’s preference for polyandry coevolves and becomes genetically coupled with the male’s sperm competitive trait, which despite its name may include non-sperm male traits (Keller and Reeve 1995; McNamara et al. 2014). These competitive traits may include the ability of the male to achieve multiple copulations with the same female (Bernasconi and Keller 2001), produce and/or transfer more sperm (Keller and Reeve 1995) or a larger ejaculate during mating, reach and fertilize the egg more successfully (Keller and Reeve 1995; Birkhead et al. 1999), displace previously stored sperm within the female’s reproductive tract (Keller and Reeve 1995; Civetta 1999; Manier et al. 2010), or prevent their own sperm in female storage from being displaced (Keller and Reeve 1995; Bernasconi and Keller 2001), which may occur by reducing female remating through chemical or physical barriers (Wolfner 1997; Baer et al. 2001; Sutter et al. 2016). Should the male competitive trait(s) and female preference for polyandry become genetically associated, positive selection for one will indirectly select for the other, which lends itself to positive feedback on both and the maintenance of polyandry (Keller and Reeve 1995).

An important assumption of the sexy-sperm hypothesis is father-to-son heritability of male traits that promote fertilization success, which explains how polyandrous females derive an indirect fitness benefit – by having sons that are more competitive. Hence, a critical prediction of this hypothesis is that polyandrous females should produce sons that are on average more successful under sperm competition than sons from monandrous females (Keller and Reeve 1995). A number of recent empirical investigations in disparate taxonomic groups, including another beetle species, have tested and found support for this prediction as well as support for the prediction that female promiscuity genes are inherited by daughters (Bernasconi and Keller 2001; Pai and Yan 2002; Klemme et al. 2008, 2014; Simmons and García-González 2008; Iyengar and Reeve 2010; McNamara et al. 2014; Egan et al. 2016). While there is mounting support for sexy-sperm processes, there are also examples for which no support was found for the evolution of sexy-sperm processes (Jennions et al. 2007; Konior et al. 2009). One potential reason for these disparate findings is that father-to-son heritability may be constrained by the genetic architecture underlying traits important in sperm competition within certain species (Evans and Simmons 2008).

The assumption of father-to-son heritability required for sexy-sperm processes to occur is contentious due to several possible theoretical constraints, all of which have some empirical support. One constraint is that there must be a response to selection by the genes in males for sperm competitiveness to evolve adaptively (Houle 1992; Keller and Reeve 1995; Pizzari and Birkhead 2002), which is only possible if the phenotypic variation in sperm competitive traits exhibits additive genetic variance and is attributable to autosomal rather than maternal (e.g., X-linkage or cytoplasmic genes derived from mitochondria) genetic effects (Radwan 1998; Birkhead et al. 1999; Morrow and Gage 2001; Simmons and Kotiaho 2007; reviewed in Evans and Simmons 2008). Further constraints include the nonadditive nature of genes (e.g., dominance, epistasis, or interactions) and trade-offs among ejaculate components (e.g., negative genetic associations due to pleiotropy or linkage disequilibrium), both of which likely impede an optimal, directional response to selection by sperm competitive traits (Moore et al. 2004; Birkhead et al. 2005; Evans and Simmons 2008).

Previous empirical work has revealed such genetic constraints in the seed beetle, *Callosobruchus maculatus*, a model system in sperm competition studies in which female multi-male mating is ubiquitous (Eady 1995). Sperm competitiveness in this species has been shown to be contingent upon parental compatibility (i.e., interactions between maternal and paternal genotypes; Wilson et al. 1997) and exhibit nonadditive genetic variation (Dowling et al. 2007b). Together these results undermine the plausibility of the response to selection necessitated by sexy-sperm processes, but whether females in this system derive such an indirect fitness benefit has never previously been tested.

Here, I test the sexy-sperm hypothesis in the seed beetle by assessing the relative sperm competitive success of sons from polyandrous females and sons of monandrous females in competitive double matings. I present new evidence that contradicts it – sons from monandrous females had significantly greater relative fertilization success than sons from polyandrous females under sperm competition. Behavioral observations of these matings were used to conduct post-hoc investigations of mechanisms for differential fertilization success between sires. My results suggest that the likely mechanism for this paternity bias is the significantly larger ejaculate produced by sons from monandrous females, which is unexpected given that these males were significantly smaller in size. Importantly, this difference in ejaculate size was only found when monandrous sons were second to mate, which is the favored role in this species which features high last male sperm precedence (Eady 1991; Hook 2017). These exciting new results give a new dimension to our understanding of the evolution of polyandry and suggest that ‘anticipatory maternal effects’ (Marshall and Uller 2007) may be another factor capable of influencing sperm allocation decisions and sperm competitive ability.

## METHODS

### CULTURING BEETLES AND DEVELOPING MATRILINES

All seed beetles used in this experiment came from an outbred culture of a single wild-type population of *C. maculatus* originally collected in southern India (Messina & Mitchell 1989) and provided by Dr. Messina of Utah State University. Once acquired, beetles were cultured in a 2-L glass jar containing approximately 750g of organic black-eyed beans (*Vigna unguiculata*, from Azure Standard in Durur, OR) in a laboratory growth chamber at Cornell University under constant conditions of 26±1°C, 10-50% RH, and with a 12:12 light/dark cycle. To establish known relationships between individuals and avoid matings between relatives, matrilines were initiated from stock populations by isolating randomly infested seeds to rear parental virgins. Once they emerged as adults from their natal seed, a randomly selected male and female were paired to mate. Mated females were assigned to a matriline using a unique identifying character and provided seeds on which to oviposit. To avoid larval competition, only seeds with single eggs glued to them were placed separately into 35mm Petri dishes. All successfully eclosed offspring were provided a unique identification number (ID) and their generation number, matriline, egg lay date, eclosion date, sex, and a qualitative estimate of their size (small, medium, or large) were recorded. All individuals in the present study were derived from 23 mated pairs across 11 matrilines and five generations. Matrilines (both maternal and paternal) were considered as random effects in statistical analyses.

### OBSERVING MATING BEHAVIOR

All experimental matings for this study were conducted between May and November of 2014 and were staged within the females’ 35mm Petri dish. Each mating pair was continuously observed until the mating was complete. Highly stereotyped and easily observed, mating in the seed beetle begins once the male has successfully inserted his aedeagus into the female’s genitalia, at which point he leans back and remains relatively motionless until the female begins to kick him. A struggle ensues until the male is fully dislodged from the female’s genitalia and the pair separates. All mating behaviors were recorded within the nearest 10s, beginning with the time the male was placed into the mating arena, the time the male leaned back (a proxy for successful genitalia insertion), the start of female kicking, and the time the pair was separated (a proxy for successful genitalia removal). *Copulation latency* was calculated as the time the male was placed in the mating arena to the time he leaned back, and *kicking latency* was calculated as the time the male leaned back until the time the female began to kick. *Kicking duration* was calculated as the time the female began to kick until the pair successfully separated, and *copulation duration* was calculated as the time the male leaned back to the time his genitalia was successfully removed. Immediately before and after each mating, all individuals were weighed twice using a Sartorious MP1601 micro-balance to the nearest ±0.1mg and resulting weights were averaged. Male ejaculate size was calculated by subtracting his post-mating weight from his pre-mating weight.

Females that were provided additional opportunities to mate were allowed to naturally vary in when they were willing to remate – either 0, 24, or 48 hours after their first mating (unless otherwise noted). Females that were unwilling to remate at these times points exhibited several resistance behaviors that prohibited successful copulation; they ran away or kicked approaching males or moved their abdomen so that the males were unable to insert their aedeagus. Some females exhibited these unreceptive behaviors at first but then eventually acquiesced and remained motionless so that the male could mount her and insert his aedeagus. For females that did not remate within the mating trial, the male was removed and she was provided with a single seed for oviposition. A second opportunity to remate was provided 24 hours later. Unreceptive females were provided their same single seed for oviposition and another opportunity was given the next day (i.e., 48 hours after the first mating). To standardize female exposure to males, pairings in which the total pairing duration exceeded 30 minutes, the male mate died prior to female remating (thus requiring her to be exposed to a different second male to remate), females mated more than twice with a male, or females remated beyond 48 hours after the first mating were removed from the study.

### MONANDROUS AND POLYANDROUS MATERNAL TREATMENTS

Female offspring derived from matrilines were randomly assigned to either a single male (‘monogamous’ treatment) or a pair of males (‘polyandrous’ treatment) for two sequential matings. Female opportunities to remate were altered over the course of the experiment, starting with more restrictive time intervals of 24- or 48-hours post-mating (or both) and then eventually expanded to include 0-, 24-, and 48-hour post-mating intervals. Importantly, all females within the same generation were given the same opportunities to remate, so mating regimes did not differ within any one generation. These differences were taken into account statistically by assigning females within a single generation a unique group number, which was considered as a random factor in the statistical model for paternity but was subsequently removed because it did not to affect paternity outcomes.

Some females were not given a choice to remate and so were under enforced monogamy (*n* = 5, treatment 4, Fig. 1A). Females that were given a choice to remate but did not do so were considered monogamous (*n* = 6, treatment 3, Fig. 1A). Of the 35 females that did successfully remate, 17 had been assigned to a single male (treatments 1 and 2, Fig. 1A) and 18 had been assigned to two different males (treatments 5 and 6, Fig. 1A). Initially, there was only one polyandrous mating treatment in the experiment, which entailed consecutive matings with two virgin males (*n* = 9, treatment 5), which are known to produce significantly larger ejaculates than previously mated males in this species (Eady 1995; Savalli & Fox 1999). Because monogamous matings necessarily entail consecutive matings with a virgin male first and a once-mated male second, a new treatment group (treatment 6) that included a virgin male as a first mate and once-mated male as a second mate was created to control for the effects of ejaculate size. Similarly, to see if previously mated males would have an effect on paternity outcomes, some females were mated twice to the same non-virgin male (*n* = 4, treatment 2).

**FIGURE 1.**
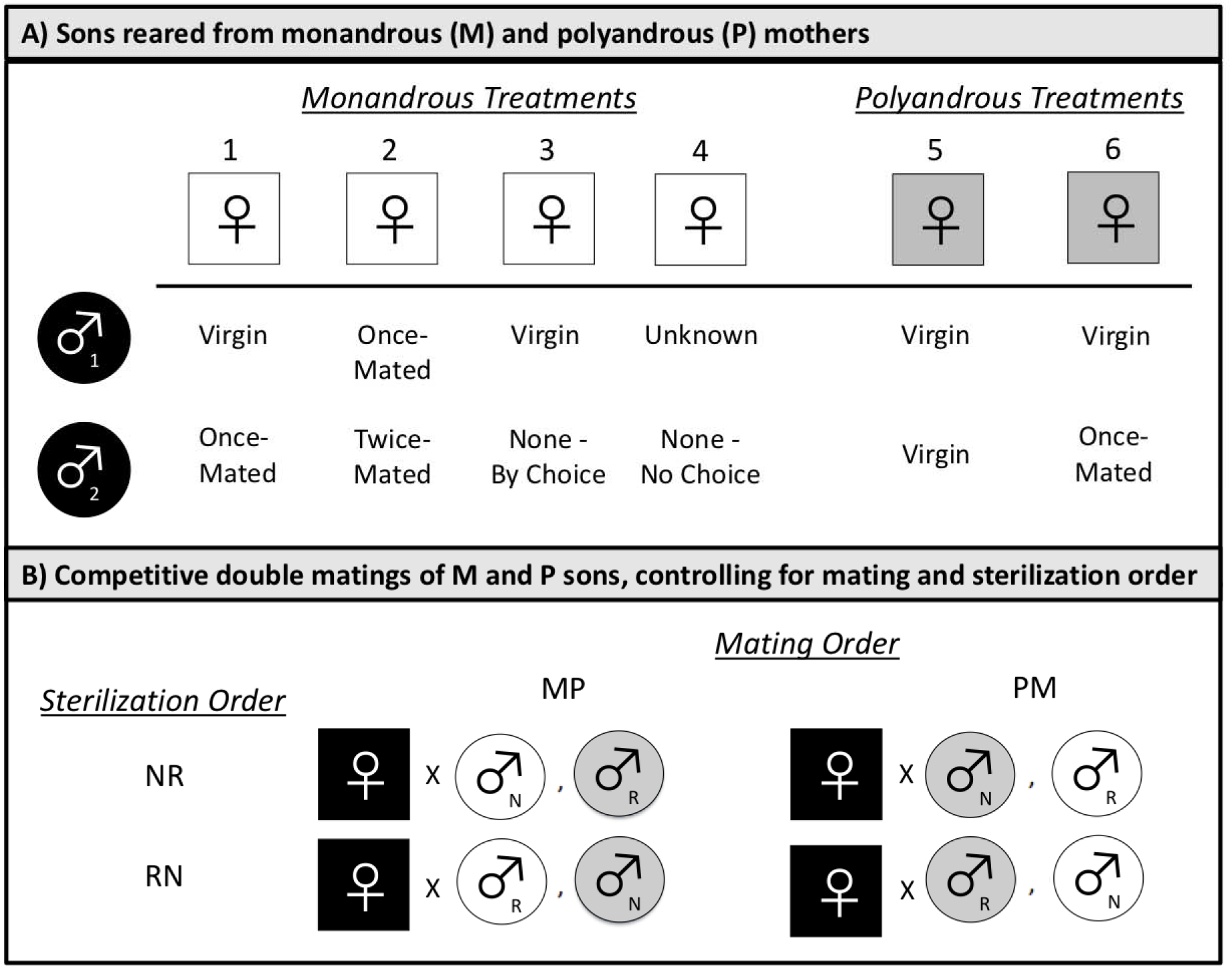
Experimental design for testing a prediction of the sexy sperm hypothesis that sons from polyandrous females have higher average fertilization success than sons from monandrous females. Matings were staged to produce a generation of virgin females (A), which were then randomly assigned to either monandrous (M) or polyandrous (P) treatments (B). Their sons (M white circles, P gray circles) served as focal virgin males in competitive double matings in which they were age- and size-matched and randomly assigned to unrelated, virgin females (black squares) (C). Paternity success of the second male to mate (P2) was determined using the sterile male technique, in which one male was irradiated (R) and the other was not (N). A balanced design was used to control for both sterilization order (RN and NR) and mating order (MP and PM).

Given all of this variation in inter-mating interval, mating frequency, number of mates, and mating status of mates, the unique treatment numbers (1-6) assigned to females with differing mating regimes was considered as a covariate in paternity analyses. Post-hoc statistical analyses showed that none of these factors had an effect on their sons’ paternity success. Thus, all maternal treatments were collapsed into “monandrous” or “polyandrous” categories. Overall there were 46 mated females, which were provided with clean seeds for oviposition. Their singly laid eggs were reared in isolation to produce the focal males to be used in competitive double matings.

### COMPETITIVE DOUBLE MATINGS OF MONANDROUS VS POLYANDROUS SONS

Matriline-derived focal females were assigned to two unrelated age- and size-matched sons from monandrous (M) and polyandrous (P) mothers for a sequential double mating. A balanced design was used to control for the mating order so that approximately half of all focal females were mated first with a polyandrous son and second with a monandrous son (PM matings, *n* = 29, Fig. 1B), and half of all females mated first with a monandrous son and second with a polyandrous son (MP, *n* = 35, Fig. 1B). All individuals used in this stage of the experiment were virgins at the time of mating, and each male was used only once. Focal females were provided with a first male mate, and those that failed to mate within 20 minutes of introduction (*n* = 2, 1.7%) were discarded from the study. After the first mating, each female was immediately provided the opportunity to mate with the second male. Females that did not remate within 20±5min were given further mating trials either 24 hours and/or 48 hours later; females that were unreceptive to remating within 48 hours after their first mating (*n* = 44, 38.3%) were discarded from the study. Given this variation in the female inter-mating interval (either 0, 24, or 48 hours after an initial mating) and its impact on paternity patterns in this species (Hook 2017), this trait was included as a fixed effect in statistical analyses of paternity. Females that successfully double mated were transferred to a new Petri dish containing clean seeds for oviposition. These females were then transferred to a new dish with clean seeds every 24 hours until their natural death, at which point their eggs were counted and scored to determine siring success for each focal male.

### PATERNITY ANALYSES

Given the high last male sperm precedence in this species (Eady 1991; Hook 2017), the proportion of offspring sired by the second male to mate (P_2_) was calculated for each double mating to compare the paternity success of monandrous and polyandrous sons when second to mate. The sterile-male technique was used to assign paternity based on egg viability. Sterile males were produced by exposure to 70 Gy of gamma radiation from a cesium source at Cornell University. These males are capable of copulating, transferring their ejaculate, and fertilizing eggs comparably to normal males but, due to genetic mutations in their sperm, eggs fertilized by their sperm fail to hatch and develop normally (Boorman & Parker 1976; Eady 1991). Eggs fertilized by sterile males are easily distinguished from eggs fertilized by normal males based on their color; whereas eggs fertilized by sterile males remain clear, eggs fertilized by normal males either turn white after hatching due to accumulation of larval frass or feature a brown dot – the head of the hatched larvae (Wilson & Hill 1989; Eady 1991). For each female, only one male mate was randomly assigned to be sterilized. A balanced design was used to control for sterilization order so that approximately half of all focal females were mated first with a normal (N) male and second with a sterile (R) male (NR matings, *n* = 31, Fig. 1B), and half of all females mated first with a sterile male and second with a normal male (RN, *n* = 33, Fig. 1B). Sterilization order was considered as a fixed effect in statistical analyses. When assigning paternity, female IDs were used so that eggs could be blindly scored as hatched or unhatched without knowledge of the sterilization order, mating order, or intermating interval. For females with an inter-mating interval >0 hours, eggs laid between matings were quantified; however, only eggs laid after the double matings were used to calculate P2 using the following formula from Boorman and Parker (1976);

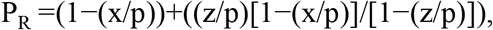

where *P_R_* represents the proportion of eggs sired by the sterile male, *x* is the proportion of eggs that hatch after a double mating, *p* is normal male fertility, and *z* is sterile male fertility.

A series of separate mating assays were conducted to determine these values within the formula. Because sterile males were irradiated on the date of the focal females’ first mating regardless of whether she remated on that date, there may be temporal effects on sterilization and fertility (Rugman-Jones and Eady 2001), particularly for sterile males assigned to the second mating role and to a focal female that delayed remating. Hence, *p* and *z* were calculated separately for both sterilization orders and each separate intermating interval (Table 1). Calculating *x* required an additional step of determining the number of brown eggs that occurred within these mating assays, since these eggs were not recorded for experimental females and were incorrectly attributed to sterile males at the time of data collection, despite having hatched. Hence, to accurately estimate the proportion of eggs that hatch after a double mating (*x*), the number of brown eggs resulting from these matings (Table 1) were summed to determine a proportion of brown eggs for each experimental focal female (based on inter-mating interval and sterilization order). This proportion was then multiplied by the number of eggs she laid after her second mating to estimate the number of brown eggs laid, which were then subtracted from the sterile male eggs and added to the normal male eggs. These final values were then used to determine *x*. Hence, all correction factors were applied on a case-by-case basis and used within the formula to estimate P_2_ for each focal female. These values were then multiplied by the total number of eggs laid after the second mating and rounded to a whole number to quantify the number of eggs fertilized by each male sire. After applying the formula, some P_2_ values were over 1.0 (*n* = 3) or under 1.0 (*n* = 1). In one of these cases, eggs that were laid between matings were used to verify that the first male mate successfully transferred sperm; this female was kept in the analysis so as not to remove important instances of paternity bias in favor of a single sire. Because it could not be ruled out for the other cases that the first or second matings were unsuccessful due to insufficient sperm transfer or male infertility, these females were excluded from the paternity analyses. Females that laid an unusually low number of eggs (<10) after their second mating (*n* = 2) were also discarded from paternity analyses. Hence, paternity was analyzed for 64 females in total.

**TABLE 1.**
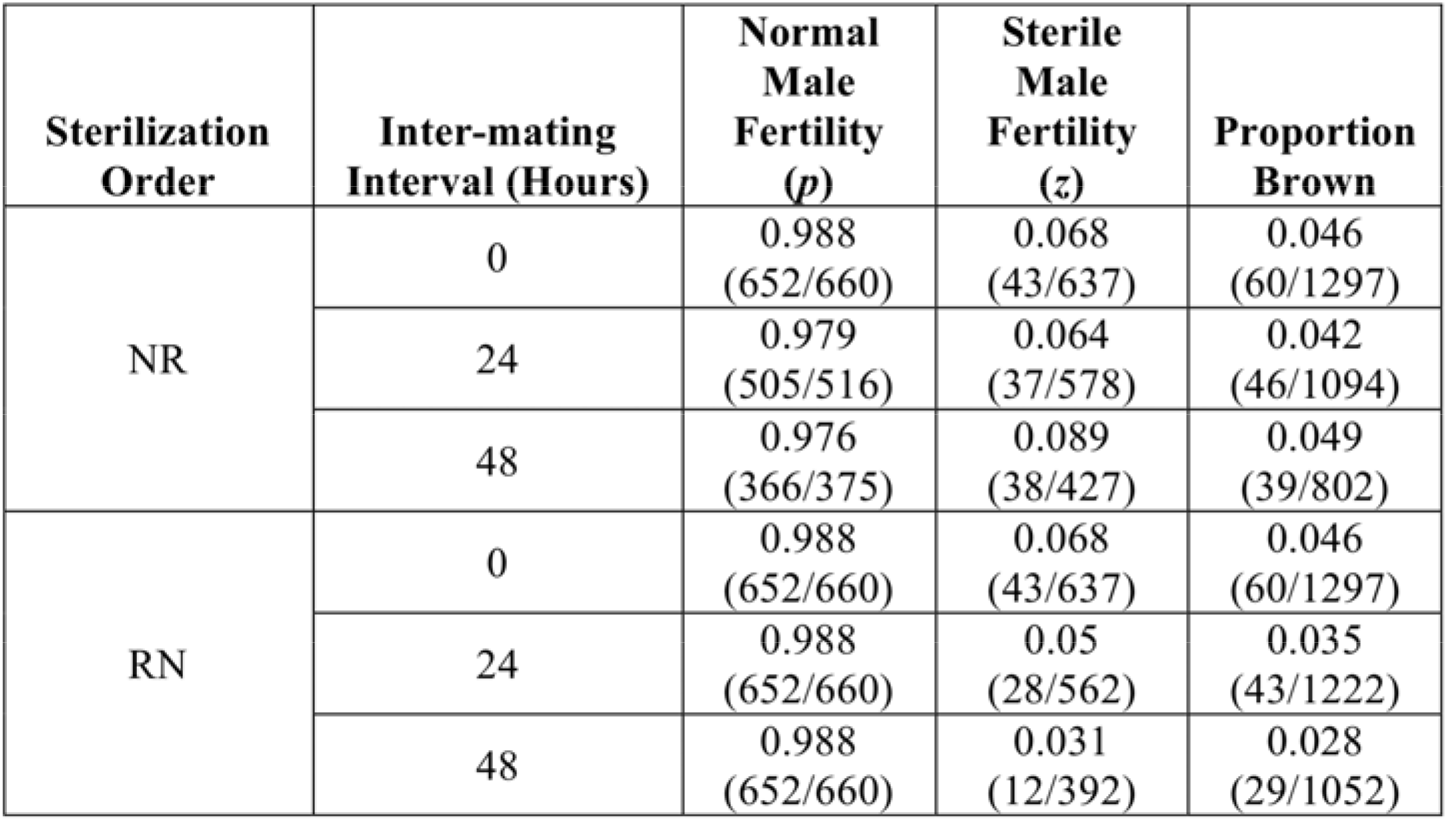
Results from mating assays to determine the proportion of eggs that successfully hatch after mating with a normal (N) male (i.e. natural fertility, p) or that erroneously hatch after mating with a sterile (R) male (i.e. incomplete sterilization, z) as well as the proportion of brown eggs laid based on a reciprocally balanced sterilization order (NR or RN) for three distinct intermating intervals (0, 24, and 48 hours after an initial mating). These proportions served as correction factors within the Boorman and Parker (1976) formula to determine paternity of offspring resulting from competitive double matings.

### POST-HOC ANALYSES FOR MECHANISMS OF PATERNITY BIAS

Post-hoc analyses were conducted to compare sons from monandrous and polyandrous females with regard to pre- and post-copulatory competition. For pre-copulatory competition, I compared the proportion of females willing to remate after a first mating and copulation latencies for both males. For post-copulatory competition, I compared kicking latencies, kicking durations, and copulation durations for both males. I further examined male body sizes, proxied by their pre-mating weights, as well as their ejaculate sizes.

### STATISTICAL ANALYSES

All statistical analyses were conducted using R version 3.1.2 (R Development Core Team 2014). All means are presented ± 1 SE.

#### PATERNITY ANALYSES

Paternity success of the second male (P_2_) was analyzed using a generalized linear mixed model (GLMM) using the glmer function from the “lme4” R package and a logit link function (Bates et al. 2015). The binomial response was the number of eggs fertilized by the second male, and the total number of eggs laid after the second mating was the binomial denominator (*n* = 64). In the initial statistical model, the residual deviance was observed to be larger than the residual degrees of freedom, which is an indication of overdispersion (Crawley 2013). An observation-level random effect (OLRE) was used as a random factor in all subsequent analyses to control for overdispersion (Harrison 2014). The experimental group (based on mating date), nested matrilines and patrilines within a generation number, and siblings (i.e., sisters for females or brothers for first and separate males) for all focal individuals were included as random factors in the initial model, but only those effects that contributed to residual variability were included in the final model. These random factors, which included female generation and an OLRE for the final GLMM, were then used in bivariate analyses for variables of interest that could potentially explain paternity bias, which included the females’ pre-mating age and weight. Because experimental males were age- and size-matched, these variables were highly collinear between males (LM male ages: F_1_,_66_ = 73.63, *P* < 0.001; LM pre-mating male weights: F_1_,_67_ = 12.64, *P* < 0.001) and so were not considered as predictors. However, second male pre-mating ages and weights were denoted at the time just before the second mating and so differed among males based on the inter-mating interval; hence, the absolute differences in pre-mating weights and ages of males were considered, as were their absolute differences in copulation latencies, kicking latencies, kicking durations, copulation durations, and ejaculate sizes. Some of these latter five predictor variables were missing for males in either the first (*n* = 15 gaps out of 320, or 4.7%) or second (*n* = 23 gaps out of 320, or 7.2%) mating role; averages were calculated for each of these variables, and these values were then used in place of missing information so that the model would not drop these males entirely from the analysis. To compare maternal differences in inter-mating intervals, mating frequency, number of mates, mating status of mates, new columns of data were created and dummy coded with a “0” if mothers did not differ and a “1” if they did differ in these traits. These comparisons, as well as maternal treatment number comparisons (1-6), were considered as predictors in the model as well. Only predictors that had a *p* value at or below 0.20 were considered for the final model. These predictors were further screened for collinearity with other significant predictors and were removed whenever collinearity was present so that only the one with greater relative significance was included in the final GLMM. Non-significant terms were dropped one at a time, and models were compared using Akaike information criterion (change in AIC < 2). Only the best fitting model is reported here. The only fixed effects in the final model included the mating order and inter-mating interval. Post-hoc comparisons were made using Tukey HSD adjustments for multiple comparisons with the “LSmeans” R package (Lenth 2016).

#### POST-HOC ANALYSES FOR MECHANISMS OF PATERNITY BIAS

The proportion of females remating at each inter-mating interval based on mating order was analyzed using a Pearson’s Chi-squared test with Yates’ continuity correction. Pre- and post-copulatory mating variables (copulation latency, kicking latency, kicking duration, and copulation duration) and male traits (pre-mating weights, ejaculate sizes) were analyzed separately using linear mixed models (LMMs) to test for significant differences between monandrous and polyandrous sons (*n* = 69). These models were run using the lmer function from the “lme4” R package (Bates et al. 2015), and some response variables were log or square root transformed to meet the model assumptions of normality and/or homogeneity of the variance. Whenever no random effects contributed significantly to residual variability in the response variables, linear models (LMs) were used instead. For LMMs involving the first mating, nested generations, matrilines, and patrilines of the female and first male and experimental group were included as random effects. For LMMs involving the second mating, nested generations, matrilines, and patrilines of all focal individuals and experimental group were included as random effects. Only those effects that significantly contributed to residual variability were included in the final models; these random effects were then used in bivariate analyses for predictors of interest. For first matings, these included female and first male ages and weights, sterilization order, and mating order. For second matings, these included female, first male, and second male ages and weights, first mating variables (e.g. copulation latency, kicking latency, kicking duration, copulation duration, ejaculate size), sterilization order, inter-mating interval, and mating order. For the male pre-mating weight LMM, male development time in the natal seed (i.e., egg laying to eclosion date) was also included as a predictor. For all LMMs, only variables with significant effects were screened for collinearity using methods outlined above before being included as covariates in the final models. Non-significant terms were dropped one at a time based on model comparisons using analyses of variance (ANOVA). Only the best fitting models are reported here.

## RESULTS

### PATERNITY ANALYSES

The proportion of offspring sired by the second male to mate (P_2_) significantly differed based on mating order and inter-mating interval (Table 2). When polyandrous sons were second to mate, P_2_ was significantly lower (polyandrous sons= 61.8 ± 3.4%, monandrous sons= 68.3 ± 3.8%; binomial GLMM: *n* = 64, *P* = 0.04; Table 3; Fig. 2). Fitted values from LSmeans for P2 were 62.9 ± 4.9% for polyandrous sons and 73.1 ± 4.4% for monandrous sons.

**TABLE 2.** Paternity results from competitive double matings in which focal females were mated to sons from polyandrous (P) and monandrous (M) mothers. Matings were reciprocally balanced for mating order (MP, PM), and all second matings occurred either 0, 24, or 48 hours after the initial mating. The proportion of offspring sired by the second male to mate (P2) was calculated separately for each inter-mating interval and mating order by dividing the total number of eggs fertilized by second males by the total number of eggs laid across females within each subgroup.

| Inter-mating Interval: | Mating Order: | Sample Size | Proportion of Offspring Sired by the Second Male to Mate ($P_2$) |
| --- | --- | --- | --- |
| 0 | MP | 13 | 417/828 = 0.50 |
|  | PM | 12 | 442/742 = 0.60 |
| 24 | MP | 15 | 373/542 = 0.69 |
|  | PM | 8 | 291/368 = 0.79 |
| 48 | MP | 7 | 143/203 = 0.70 |
|  | PM | 9 | 182/252 = 0.72 |

**TABLE 3.** Fixed effects from generalized linear mixed model examining the effects of mating order, inter-mating interval, and sterilization order on the proportion of eggs fertilized by the second male to mate (P2). Female generation and an observation level random effect (OLRE) were the only random factors included in the final model. Significant terms are bolded.

|  | Paternity Success of Second Male |  |  |
| --- | --- | --- | --- |
|  | N = 64 |  |  |
| Model Term | Beta ± SE | z | p |
| Intercept | 0.09 ± 0.25 | | |
| Mating Order (PM) | 0.47 ± 0.22 | 2.10 | **0.0357** |
| Inter-mating Interval (24h) | 0.82 ± 0.26 | 3.21 | **0.0013** |
| Inter-mating Interval (48h) | 0.51 ± 0.29 | 1.76 | 0.0779 |

**FIGURE 2.**
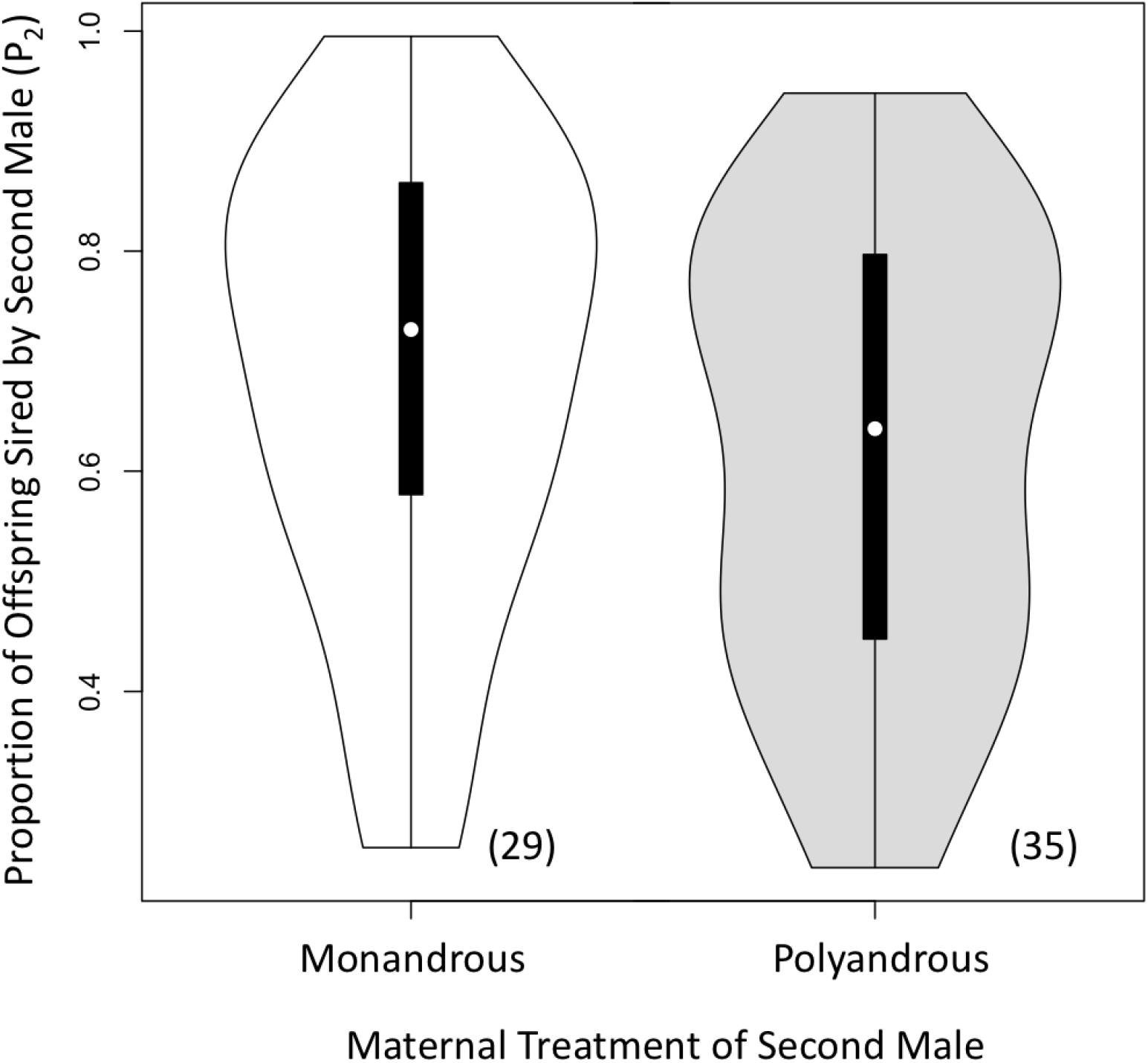
The proportion of offspring sired by the second male to mate (P_2_) in competitive double matings based on the maternal treatment of males in the second mating role. Matings in which the maternal treatment of males in the second mating role were polyandrous (P) are shaded in gray. Sons from monandrous (M) females (shaded in white) sired significantly more offspring when second to mate than sons from polyandrous females, which is the opposite of what is predicted by the sexy-sperm hypothesis. Violin plots show P_2_ (median is white dot, interquartile ranges are black bars, and width represents the probability density). Sample sizes are written in parentheses.

### POST-HOC ANALYSES FOR MECHANISMS OF PATERNITY BIAS

The proportion of females remating did not significantly differ between females that mated first with a polyandrous or a monandrous son for each separate inter-mating interval (Table 4). Moreover, polyandrous sons did not significantly differ from monandrous sons in their competitive mating behaviors (e.g., copulation latency, kicking latency, kicking duration, or copulation duration) when either first or second to mate (Table 5). When in the first male mating role, polyandrous sons showed a pattern of mating for a slightly longer duration of time, which was only borderline significant (Table 5). When first to mate, polyandrous and monandrous sons did not differ in the size of their ejaculates; however, when second to mate, polyandrous sons produced significantly smaller ejaculates than monandrous sons (Table 5, Fig. 3). Further analyses were conducted to see if these polyandrous sons were smaller in size, but I found the opposite to be the case – polyandrous sons weighed significantly more prior to mating than monandrous sons (poly sons = 3.86 ± 0.08mg; mono sons= 3.66 ± 0.07mg; LM: n = 140, *P* < 0.001; Fig. 4).

**TABLE 4.** The proportion of females that remated 0, 24, or 48 hours after their first mating as a function of whether their first mating partner was a son from a monandrous mother or a son from a polyandrous mother.

| Proportion of Females that Remated |  |  |  |
| --- | --- | --- | --- |
| Maternal Treatment of First Mate | Inter-mating Interval (hours) |  |  |
|  | 0 | 24 | 48 |
| Monandrous | 0.23 (13/57) | 0.34 (15/44) | 0.28 (08/29) |
| Polyandrous | 0.23 (13/56) | 0.23 (10/43) | 0.30 (10/33) |
| | $\chi^2=0.00, p = 1.00$ | $\chi^2=0.70, p = 0.38$ | $\chi^2=0.00, p = 1.00$ |

**TABLE 5.**
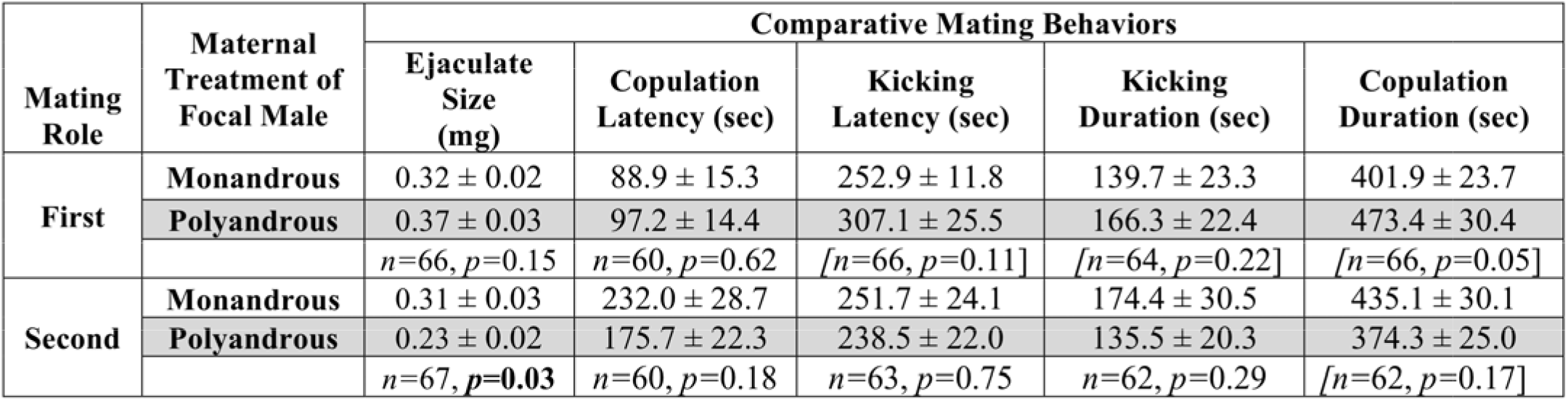
Mean (±SE) mating behavioral traits of sons from polyandrous and monandrous females based on their mating role (first or second to mate) within the competitive double matings. Sons from polyandrous females (shaded in gray) did not significantly differ in their mating behaviors, but they did produce a significantly smaller ejaculate when second to mate than sons from monandrous females. Statistics are based linear models or linear mixed models (denoted in brackets) for each separate response variable. Significant terms are bolded.

**FIGURE 3.**
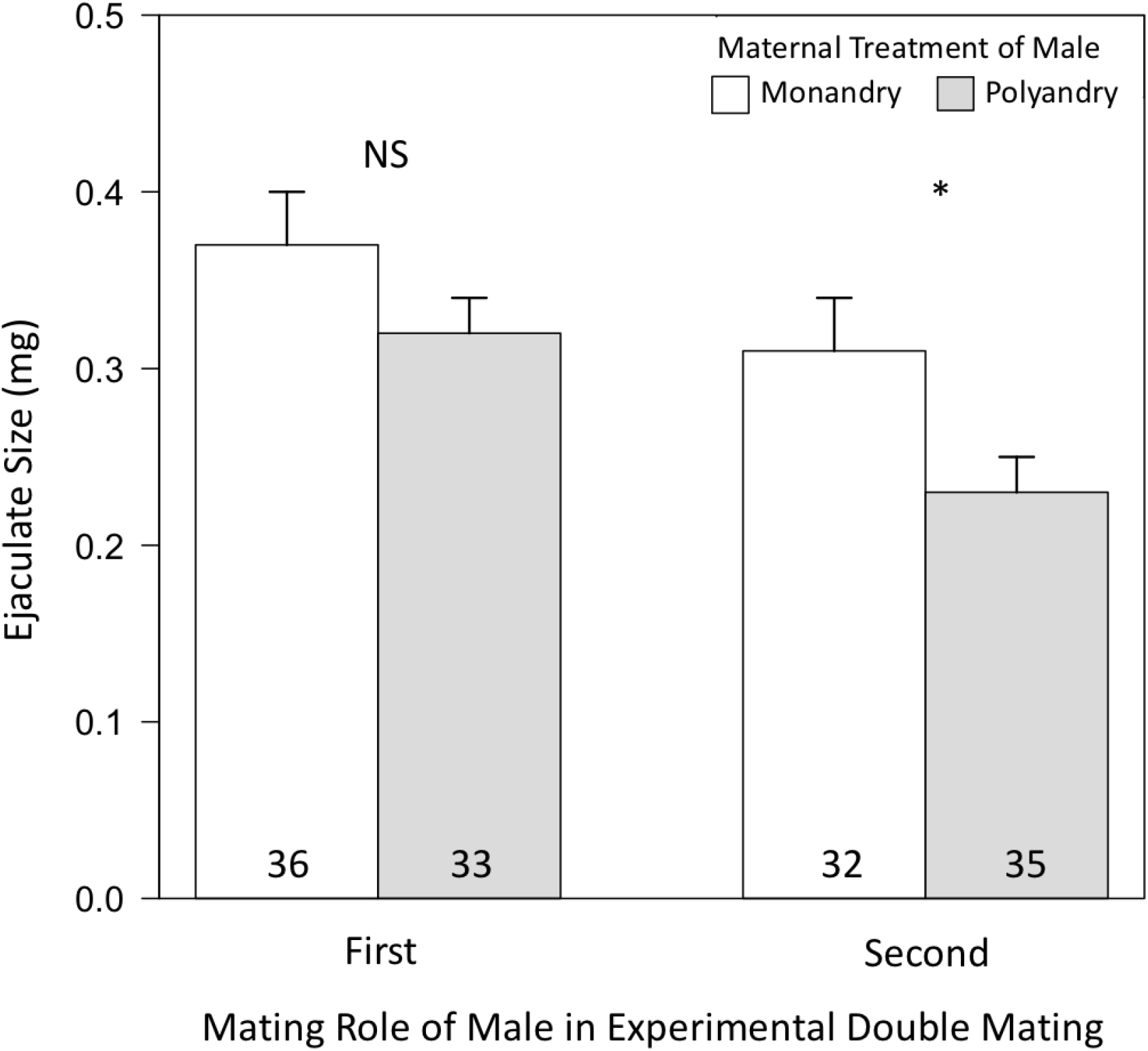
Mean (±SE) ejaculate sizes (mg) of sons from polyandrous and monandrous females based on their mating role (first or second to mate) within the competitive double matings. When first to mate, sons from monandrous mothers (white bars) did not produce a different sized ejaculate than sons from polyandrous mothers (gray bars). However, when second to mate, sons from monandrous mothers produced a significantly larger ejaculate than polyandrous sons, which is the opposite of what is predicted by the sexy-sperm hypothesis. Sample sizes are written in bars.

**FIGURE 4.**
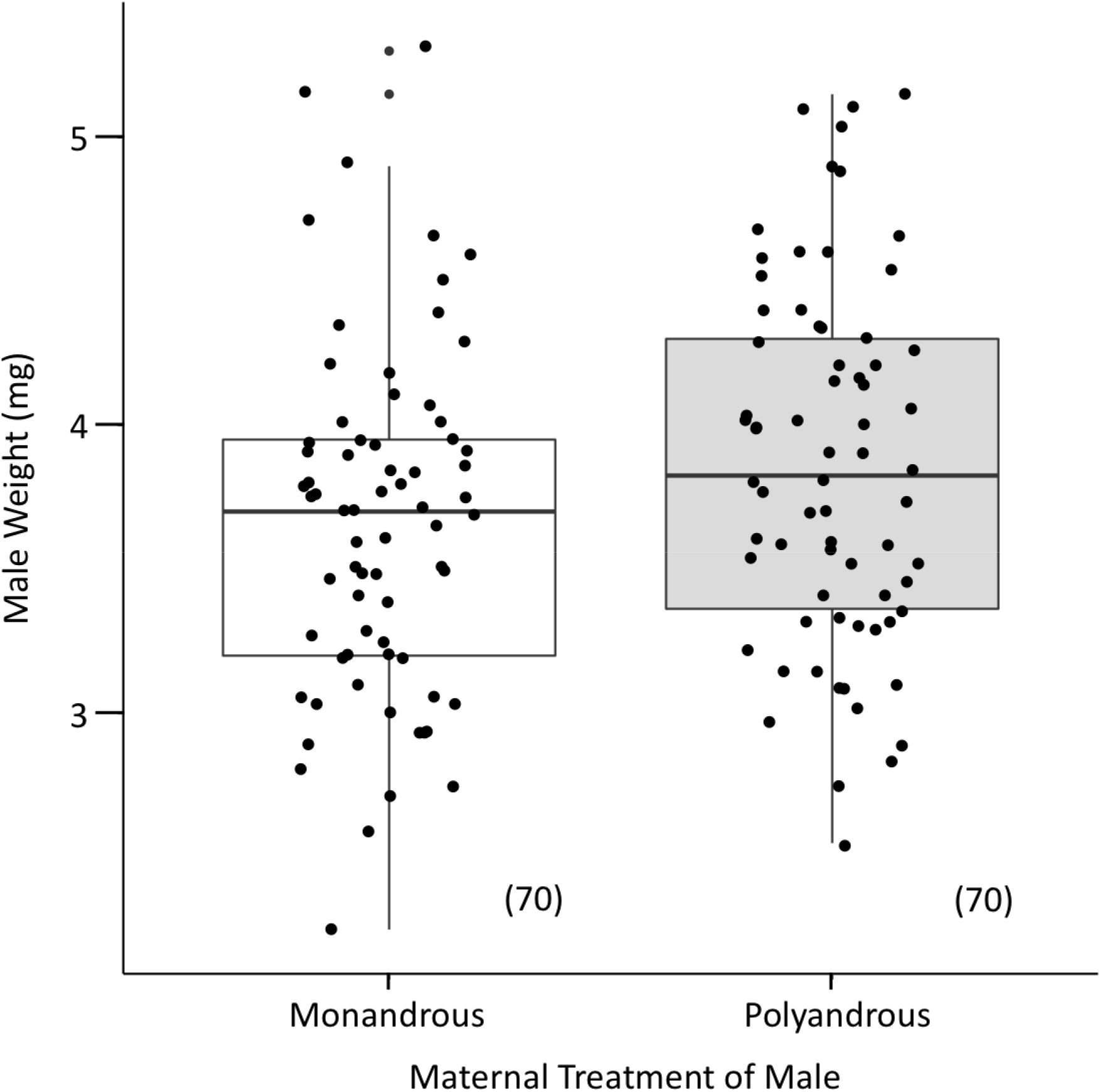
The pre-mating body weights of focal males based on maternal treatment. Controlling for age, sons from polyandrous females (gray) weighed significantly more than sons from monandrous females (white). Box-plots show body weight (medians, interquartile ranges, whiskers) and are overlaid with raw data points. Sample sizes are written in parentheses.

## DISCUSSION

The purpose of this study was to empirically test the sexy-sperm hypothesis in the seed beetle, *C. maculatus*, which has been shown to have genetic constraints that limit the likelihood of father-to-son heritability required for sexy-sperm processes to occur. Unexpectedly, the results of the present study contradict the sexy-sperm hypothesis; monandrous sons had significantly greater relative paternity success than polyandrous sons. Intriguingly, monandrous sons produced significantly larger ejaculates than polyandrous sons, despite being significantly smaller in size, but only when second to mate. This study is the first, to my knowledge, to provide empirical evidence for post-copulatory processes favoring sons from monandrous females.

Despite the fact that several empirical studies across disparate taxonomic groups have provided support for the sexy-sperm hypothesis, the results from the present study indicate that sexy-sperm processes are unlikely to occur in the seed beetle. This result is perhaps unsurprising given the underlying genetic architecture for sperm competitive traits revealed through previous studies in this species (Wilson et al. 1997; Dowling et al. 2007a). It’s possible that a directional and optimal response to selection by sperm competitive traits in this species is prohibited by nonadditive genetic variation (Pizzari & Birkhead 2002; Evans & Simmons 2008), as has been found in the fruit fly (Clark et al. 1999; Clark 2002). Yet another possibility is that the seed beetle’s XY sex chromosome system makes it less conducive to sexual selection (Smith & Brower 1974; Reeve & Pfennig 2003) or that coevolution of male fertilizing efficiency and female polyandry is restricted in this system due to either mating order effects on sperm precedence (Bocedi & Reid 2015) or sex linkage (Kirkpatrick & Hall 2004). Alternatively, sexy-sperm processes may not evolve when the costs to females from the evolution of sperm competitive traits and genetic linkage exceed the benefits for polyandry and conferral of these traits on their sons (Pizzari & Birkhead 2002). Yet another possibility is that the design of the present study may lack the power to detect heritable sperm ability and so may not allow the effects of polyandry over multiple generations to be realized.

A surprising result from this study was that monandrous sons had significantly higher paternity success than polyandrous sons. While several previous empirical studies have demonstrated support for the prediction that polyandrous sons fare better in competition, few have revealed a competitive advantage for monandrous sons. In the field cricket, monogamous sons matured more rapidly and were favored in pre-mating bias because they were more likely to win an encounter in direct mate competition against size-matched polyandrous sons (Jennions et al. 2007). In the red flour beetle, monogamous sons outcompeted polyandrous sons when mating in the first male role, but the opposite pattern was observed when they were second to mate (Bernasconi & Keller 2001). Moreover, female fruit flies mated singly (i.e., in a non-competitive context) to males from monogamous lines produced offspring at a faster rate and produced more surviving offspring than females mated to males from polyandrous lines (Pitnick et al. 2001). The present study is the first to demonstrate an advantage to monandrous sons in post-copulatory processes within a competitive context.

One possible explanation for the fertilization bias towards monandrous sons is that these males were in better condition because their mothers or fathers were of higher quality. Both scenarios are unlikely for several reasons. First, monandrous mothers did not weigh more and were not younger or more fecund than polyandrous mothers, which suggests similar body conditions. It’s possible that condition differences went undetected if harm was incurred internally through mating itself. Males in this species have elongated spines on their aedeagus that cause damage to the female reproductive tract during mating (Crudgington and Siva-Jothy 2000; Hotzy and Arnqvist 2009; Hotzy et al. 2012). If physiological harm and mating costs are incurred cumulatively, then females that only mated once may have been in better condition than females that mated multiply. This is unlikely, however, given that monandrous mothers did not differ from polyandrous mothers in their lifespans or lifetime fecundities. Furthermore, the experimental design of this study allowing females to self-select for remating could have exacerbated the likelihood that females unwilling to remate had a higher quality first male mate (‘intrinsic male quality’ hypothesis; Jennions & Petrie 2000) than females willing to remate to ‘trade up’ low quality first mates (Halliday & Arnold 1987). While it cannot be ruled out that females based their remating behavior on the intrinsic quality of their first mates, it is unlikely that females mated with males of differing qualities overall because we expect polyandrous females to remate with males of higher quality and only those females that did remate were included within the study. Together these results do not suggest there were condition differences in the parents of focal males.

Another possibility is that monandrous sons were in better condition because their mothers allocated more resources to them. It cannot be ruled out that monandrous mothers compensated for a single mate or mating by maternally allocating more to their sons, which may include material benefits derived from the ejaculate (Simmons 2005; Edvardsson 2007). Were this to be the case, we might expect these females to be constrained to lay fewer eggs (Fox and Mosseau 1998), but this explanation is not supported as mothers did not differ in the overall quantity of eggs they laid. Other adaptive maternal effects that cannot be ruled out is that monandrous mothers varied the sizes or contents of eggs, as has been demonstrated in another seed beetle (Fox and Mosseau 1998), or that they oviposited on higher quality (i.e., more nutritious) host seeds (Mosseau and Fox 1998). If we may indirectly infer maternal investment in eggs from their sons’ body size, however, then results are contrary to what we would predict since monandrous sons weighed significantly less than polyandrous sons, not more. It seems equally unlikely that developmental conditions that aid male offspring in being more competitive do not scale linearly with their body weight and so went undetected.

Other possible explanations for paternity bias in favor of monandrous sons involve the focal females and focal males in the competitive double matings. Because all copulations were directly observed in this study, potential mechanisms for the paternity bias favoring monandrous sons could be investigated, but none of these behavioral traits were found to differ between monandrous and polyandrous sons. Moreover, focal females did not differ in the number of eggs laid after the second mating (used to calculate P_2_) or the amount of weight they gained after mating based on the mating order.

The most likely mechanism for paternity bias favoring monandrous sons is that they produced significantly larger ejaculates when second to mate, which corroborates a previous finding in this species that sperm precedence is in part determined by the number of sperm inseminated by the second (but not first) male (Eady 1995). However, this result defies sexy-sperm predictions and is perplexing given the significantly smaller pre-mating weights of these males, which typically produce smaller ejaculates in this species (K. Hook, unpublished data). If males in this species are capable of detecting female mating status, as has been shown in other invertebrates (Wedell 1992; Siva-Jothy & Stutt 2003), then this result suggests that these males may be making an adaptive decision to increase mating investment given the high last (second) male sperm precedence observed within this system. Future work is needed, however, to verify that males are capable of assessing female mating status to support that such decision-making for males is indeed adaptive. A non-mutually exclusive hypothesis is that females, which have been demonstrated to receive a hydration benefit through the ejaculate (Edvardsson 2007), prefer males that produce larger ejaculates and are capable of biasing paternity in favor of males that provide them (Eberhard 1996). Further experiments are needed to demonstrate this form of cryptic female choice, however.

A cross-population study found that ejaculate size in the seed beetle exhibits an additive genetic autosomal pattern of inheritance (Savalli et al. 2000), which would allow for an optimal, directional response to selection by sperm competitive traits. Under sperm competition theory, males that anticipate future competitors ought to reduce their ejaculate contribution, though this should vary among species based on occupied mating roles and sperm precedence patterns (Parker 1970; Simmons 2001; Parker & Pizzari 2010). When in the second male mating role, monandrous sons in the present study produced significantly larger ejaculates than polyandrous sons, which supports a prediction of a previous sperm competition model that male ejaculate weights should increase with low female remating rates but decrease with high female remating rates (Parker & Ball 2005). It’s possible that this result reveals a male reproductive strategy to maximize fitness by optimally adjusting ejaculate size based on sperm competition risk and partitioning their ejaculate among successive mates (Dewsbury 1982; Pitnick & Markow 1994). The larval environment in another invertebrate system has been found to affect male development and reproductive allocations (Gage 1995), but given that larval environments did not vary for the males in the present study, it is unclear how males would be able to anticipate future mating opportunities. One possibility is that mothers can provide information to their offspring about socio-sexual experience, population density, or sex ratio. Given that the seed beetle’s mating system is akin to a scramble competition with protandrous and asynchronous emergences, it may be beneficial for mothers to transmit this information to their offspring so they may plastically adjust their development or eclosion timing and, hence, allocation to their size and/or gamete production and fecundity based on fluctuating sex ratios. An important next step is to look for direct evidence that females are capable of transmitting such a signal and the mechanism through which they do so.

## DATA ACCESSIBILITY STATEMENT

Data available from the Dryad Digital Repository: [WEB LINK TO BE INCLUDED UPON ACCEPTANCE].

## AUTHORSHIP CONTRIBUTIONS

KAH designed the research, analyzed the data, and wrote the manuscript.

## ACKNOWLEDGMENTS

I thank Janis Dickinson, Hudson Kern Reeve, Linda Rayor, Mike Webster, and Heidi Fisher as well as colleagues for critical discussion and helpful comments; Frank Messina for providing the stock population of beetles, extra oviposition substrate, and advice on culture maintenance; Ken Kemphues for help with use of the irradiator; Pat Sullivan and Andy Clark for their input on experimental design; Cornell undergraduates Abigail Giancola and Hannah Yoo for beetle culturing assistance; Lynn Johnson, Fran?oise Vermeylen, and Ailene Ettinger for statistical advice; Lars Washburn for use of a microbalance; and family and friends for their support. This research was supported by grants from the Department of Neurobiology and Behavior at Cornell University, ATHENA Fund and the Linda and Samuel Kramer Graduate Student Fellowship at the Cornell Lab of Ornithology, the Cornell Sigma Xi Student Research Grant, the American Museum of Natural History, and a National Science Foundation Graduate Research Fellowship (Cornell NSF Grant DGE-1144153).

